# Hyperactivation of L-lactate oxidase by liquid-liquid phase separation

**DOI:** 10.1101/2020.12.08.416958

**Authors:** Tomoto Ura, Ako Kagawa, Hiromasa Yagi, Naoya Tochio, Takanori Kigawa, Tsutomu Mikawa, Kentaro Shiraki

## Abstract

Liquid droplets formed by liquid-liquid phase separation are attracting attention as functional states of proteins in living cells. Liquid droplets are thought to activate enzymatic reactions by assembling the required molecules. Thus, liquid droplets usually increase the affinity of an enzyme to its substrates, leading to decreased *K*_M_ values. In this study, we demonstrate a new mechanism of enzyme activation in the droplets using Llactate oxidase (LOX). In the presence of poly-L-lysine (PLL), LOX formed droplets with diameters of hundreds of nanometers to tens of micrometers, stabilized by electrostatic interaction. The enzyme activity of LOX in the droplets was significantly enhanced by a fourfold decrease in *K*_M_ and a tenfold increase in *k*_cat_. To our knowledge, this represents the first report for increasing *k*_cat_ by the formation of the liquid droplet. Interestingly, the conformation of LOX changed in the liquid droplet, probably leading to increased *k*_cat_ value. Understanding enzyme activation in the droplets provides essential information about enzyme function in living cells in addition to biotechnology applications.

## 1. Introduction

Liquid-liquid phase separation has attracted attention as a functional state of proteins in living cells.^1^ The liquid droplets formed by liquid-liquid phase separation compartmentalize biological functions in a crowded cellular environment. The reversible assembly of necessary molecules occurs due to multivalent interactions of intrinsically disordered proteins (IDPs) or nucleic acids.^2^ It has recently been reported that droplets control some intracellular reactions, such as the immune response,^3^ transcription/translation,^4,5^ and carbon dioxide fixation in cyanobacteria.^6^ Because droplets are present in prokaryotes^7,8^ and eukaryotes, they are essential in controlling the biological reactions necessary for life.

The liquid droplets can activate enzyme reactions.^1,2,9,10^ Indeed, enzymes interact favorably with IDPs and nucleic acids. For example, RubisCO, an enzyme of more than 500 kDa for carbon dioxide fixation, functions by forming droplets with a small IDP.^6^ Also, multienzyme assemblies that activate multistep reactions exhibit liquid-like properties,^11^ and they require IDP domains.^12^ Furthermore, many metabolic enzymes act as RNA-binding proteins,^13^ which often form droplets. Some *in vitro* studies have shown that enzymes, such as ribozyme,^14^ kinase,^15^ multienzyme complexes,^16^ RNA polymerase and ribosomes,^17^ are activated in droplets. Thus, many kinds of enzymes may form liquid droplets with IDPs or RNAs, leading to changes in enzyme activity within living cells. Although various mechanisms of enzyme activation in droplets have been proposed,^9,10^ the only demonstrated mechanism to date has been compartmentalization.^14^ Thus, we hypothesize that enzyme conformational changes play an important role in activating enzymes within liquid droplets due to a crowded environment.

In this study, we investigated the relationship between enzyme kinetics and liquid droplets using L-lactate oxidase (LOX) (Fig. 1). The PLL was used as a model IDP because the sequence of IDP is enriched in charged residues, especially lysine. The liquid droplet of LOX with PLL increased the enzyme activity one order of magnitude or higher than the dispersion state in a buffer solution. The activation of the enzyme in liquid droplets resulted in an increased catalytic constant (*k*_cat_) of LOX and decreased Michaelis constant (*K*_M_). The increase of the *k*_cat_ was considered to result from the conformational change of LOX in the liquid droplet. The enzyme hyperactivation in liquid droplets has an important implication not only as a control mechanism of the enzyme reaction in living cells but also in industrial applications.

**Figure 1.**
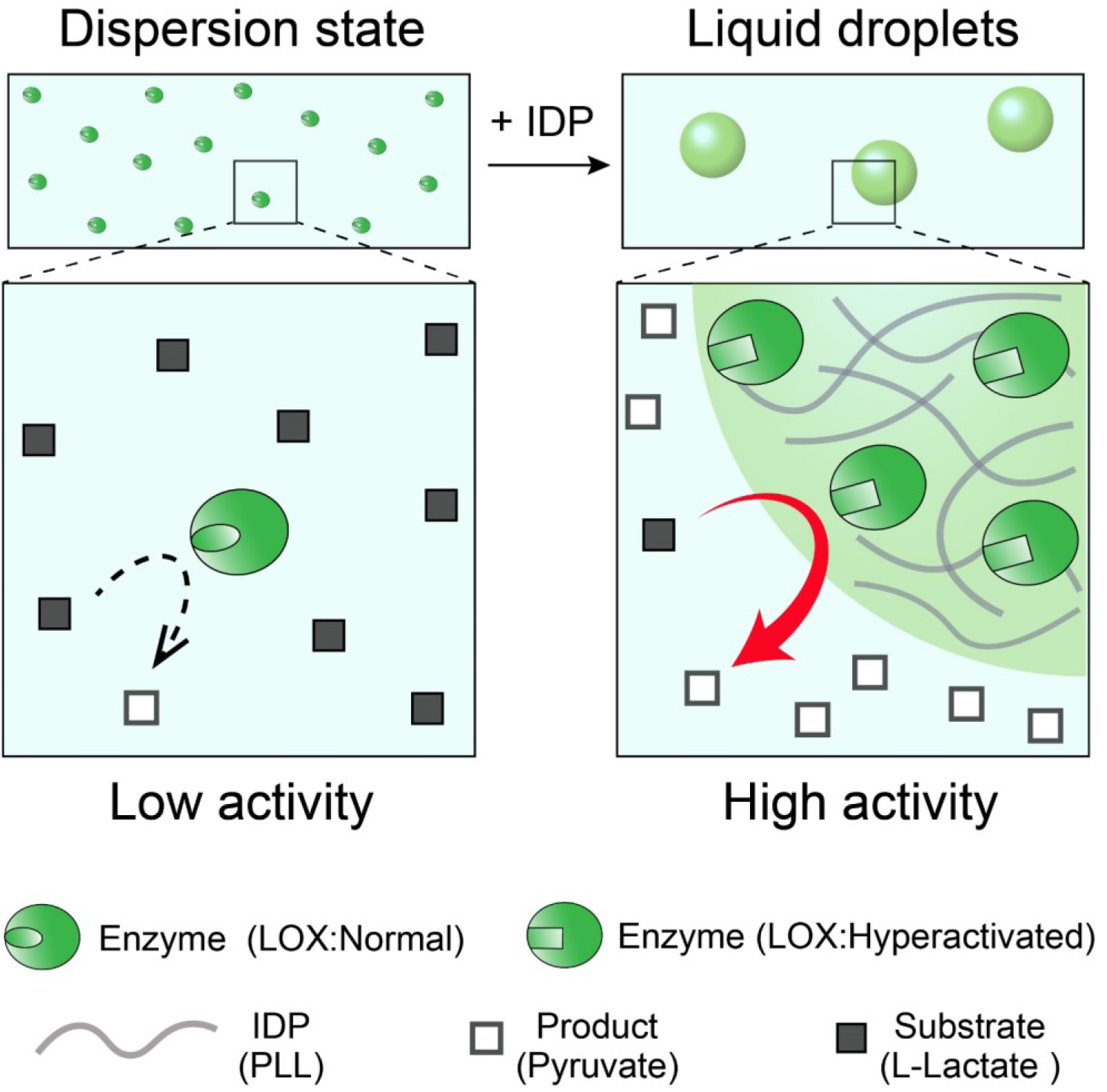
Schematic diagram of enzyme reactions in dispersion state or liquid droplets. Poly-L-lysine (PLL) mimics intrinsically disordered protein (IDP).

## 2. Materials and Methods

### 2.1. Materials

Poly-L-lysine hydrobromide (molecular weight, MW, 4,000–15,000 Da), poly-(D,L)-lysine hydrobromide (MW 25,000–40,000 Da), and 2,6-dichloroindophenol sodium salt hydrate (DCIP) were purchased from Sigma-Aldrich Co. (St Louis, MO, USA). Sodium chloride was obtained from Kanto Chemical Co., Inc. (Tokyo, Japan). Tris(hydroxymethyl)aminomethane was obtained from Nacalai Tesque (Kyoto, Japan), and 2-Morpholinoethanesulfonic acid monohydrate was purchased from Dojindo Lab (Kumamoto, Japan). Rhodamine B isothiocyanate (RBITC) was obtained from Santa Cruz Biotechnology, Inc. (Dallas, TX, USA). LOX was prepared as described previously^18^.

### 2.2. Fluorescent labeling of PLL

PLL labeled with amine-reactive dye RBITC (excitation/emission: 555/580 nm) was prepared per the manufacturer’s instructions. Briefly, a solution of RBITC (1.77 mM) in DMSO (50 μL) was quickly added to a stirred solution of 20 mM PLL in 20 mM EPPS (950 μL; pH 8.5) at room temperature. After the reaction mixture was gently stirred for 1.5 h, 200 mM Tris-HCl (100 μL; pH 8.5) was added. The PLL-dye conjugates were purified by filtration through Amicon Ultra-0.5 mL centrifugal filters with a molecular weight cutoff (MWCO) of 3 kDa (Millipore Sigma). The final concentration of PLL was determined with a bicinchoninic acid (BCA) assay. The number of RBITC molecules conjugated to each LOX enzyme in 10 mM Tris-HCl (pH 8.0) was determined from the absorbance at 556 nm using the molar absorption coefficient ε_556_ = 87,000 M^−1^cm^−1^. The number of dye molecules per PLL molecule was 0.2.

### 2.3. Enzyme assays

An enzyme solution containing 0–5 μM LOX and 0–10 mM PLL in 20 mM Tris-HCl and 20 mM MES was prepared and left standing for 20 minutes. A 260 μL aliquot of enzyme solution was mixed with a 30 μL aliquot of substrate solution containing 0–80 mM L-lactic acid and 15 mM DCIP solution. The initial reaction velocities (*v0*) were determined from the slope of the initial decrease in the absorbance at 600 nm using a JASCO spectrophotometer V-550 (JASCO Co., Ltd, Tokyo, Japan). The normalized enzyme activity was defined as the ratio of *v*_*0*_ in the presence of droplets to *v*_*0*_ in the absence of droplets.

### 2.4. Dynamic light scattering

Dynamic light scattering (DLS) experiments were performed using a Zetasizer Nano ZS light scattering photometer (Malvern Instruments, Worcestershire, UK) equipped with a 4 mW He–Ne ion laser (λ = 633 nm). For determination of the sizes of the enzyme with polyelectrolytes, solutions containing 20 nM LOX, 0–1 mM PLL, 20 mM Tris-HCl, and 20 mM MES were placed in a 1 cm path length disposable cuvette, and DLS measurements were performed at 25°C at a detection angle of 173°. The viscosity of the solutions was approximated by water (η = 0.87 cP). All results are presented as the mean values of three independent experiments.

### 2.5. Optical microscopy

All images were recorded from an all-in-one fluorescence microscope BZ-X710 (KEYENCE, Osaka, Japan). Aliquots (100 μL) of the samples were placed in an ultra-low attachment 96-well plate (Corning, NY, USA). All images were prepared for presentation in BZ-X Analyzer (KEYENCE).

### 2.6. Isothermal Titration Calorimetry (ITC)

ITC experiments were performed on a Microcal Auto-iTC200 calorimeter (Malvern Instruments). The experiments consisted of a series of 0.2 μL injections of 4 mM PLL into 200 μL of 200 μM LOX solution or 1 mM L-lactic acid in the thermostatic cell with an initial delay of 60 s, a 0.4 s duration of injection, and a spacing of 120 s between injections. In all cases, the samples were dialyzed in the same buffer of 20 mM Tris-HCl and 20 mM MES (pH 8) to minimize the interference of mixing and dilution heat signals.

### 2.7 Hydrogen-1 nuclear magnetic resonance spectroscopic analysis

Hydrogen-1 (^1^H) nuclear magnetic resonance (NMR) spectra were recorded in 20 mM Tris-HCl buffer at pH 8 (adjusted with HCl). Experiments were performed on a Bruker BioSpin Avance III 700 MHz NMR spectrometer at 25°C.

### 2.8. Circular dichroism

Circular dichroism (CD) experiments were performed in a 1 cm path-length quartz cuvette using a spectropolarimeter (J-720 W; JASCO Co., Ltd). The enzyme solution containing 1 μM LOX and 5 mM Tris-HCl buffer (pH 7.0) was incubated with 1 mM PLL or 1 mM Poly-(D,L)-lysine at 25°C for 20 minutes before measurement. The CD spectra of the samples were corrected by subtracting the corresponding spectra of buffers.

## 3. Results

### 3.1 Liquid droplets of LOX and PLL

LOX has an isoelectric point around pH 6; hence, it is negatively charged under physiological pH. Because PLL has an isoelectric point of pH 10 and a disordered structure, mimicking IDP, PLL was selected as the pair polymer of LOX. Figure 2 shows the bright-field and fluorescence microscopy micrographs of the sample containing 5 μM LOX and 1 mM PLL. To observe LOX and PLL independently, PLL was chemically modified with RBITC (red), whereas intrinsic flavin mononucleotide (green) was used for LOX. Bright-field microscopy showed droplets with spherical structures and diameters of ten micrometers or less (Fig. 2A), indicating the typical appearance of liquid droplets. Fluorescence microscopy revealed that the droplets uniformly contained both LOX and PLL molecules, as judged by green and red fluorescence. Furthermore, LOX-PLL droplets coalesced in approximately 10 s (Fig. 2B), indicating that the LOX-PLL droplets have liquid-like fluidity.

**Figure 2.**
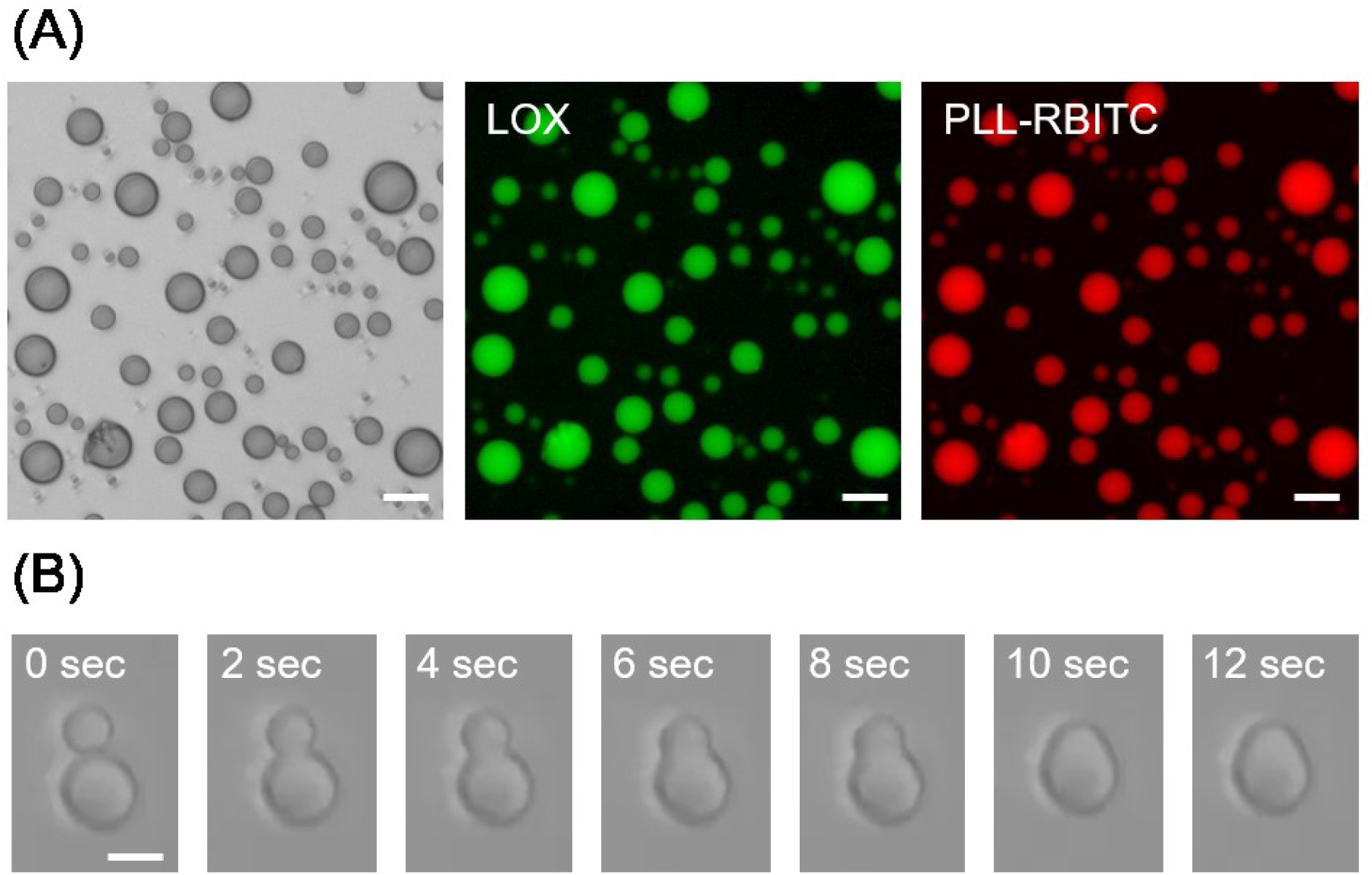
Formation of LOX-PLL droplets. (A) Bright-field microscopic images of droplets (left) and fluorescent microscopic images of LOX (middle) and PLL-RBITC (right). The solution contained 5 μM LOX, 1 mM PLL, 20 mM Tris HCl, and 20 mM MES (pH 8). Scale bar, 20 μm. (B) Bright-field microscopic images of LOX-PLL droplets. The solution contained 5 μM LOX, 1 mM PLL, 20 mM Tris HCl, 20 mM MES (pH 8). Scale bar, 10 μm.

### 3.2 LOX hyperactivation in LOX-PLL droplets

To investigate the effect of liquid droplets on enzyme activity, we measured the bright-field microscopic images and enzyme activity of LOX under various conditions. In LOX-PLL mixtures containing 5 μM LOX with 1–10 mM PLL (Fig. 3A), the LOX-PLL droplets were observed at PLL concentrations above 100 μM. Spherical droplets formed in the absence of NaCl, whereas the droplets disappeared at above 200 mM NaCl (Fig. 3B). Thus, LOX-PLL droplets were dependent on NaCl concentration, indicating that the electrostatic interaction between PLL and LOX plays an important role in droplet formation. Figure 3C shows the microscopic images of LOX-PLL droplets at different pH values. At pH 5, only amorphous aggregates were observed. Both amorphous aggregates and liquid droplets were observed at pH 6 and 7, whereas only liquid droplets were observed at pH 8 and 9. These results also demonstrate that the electrostatic interaction between LOX and PLL plays a vital role in forming liquid droplets because LOX has an isoelectric point around pH 6.

**Figure 3.**
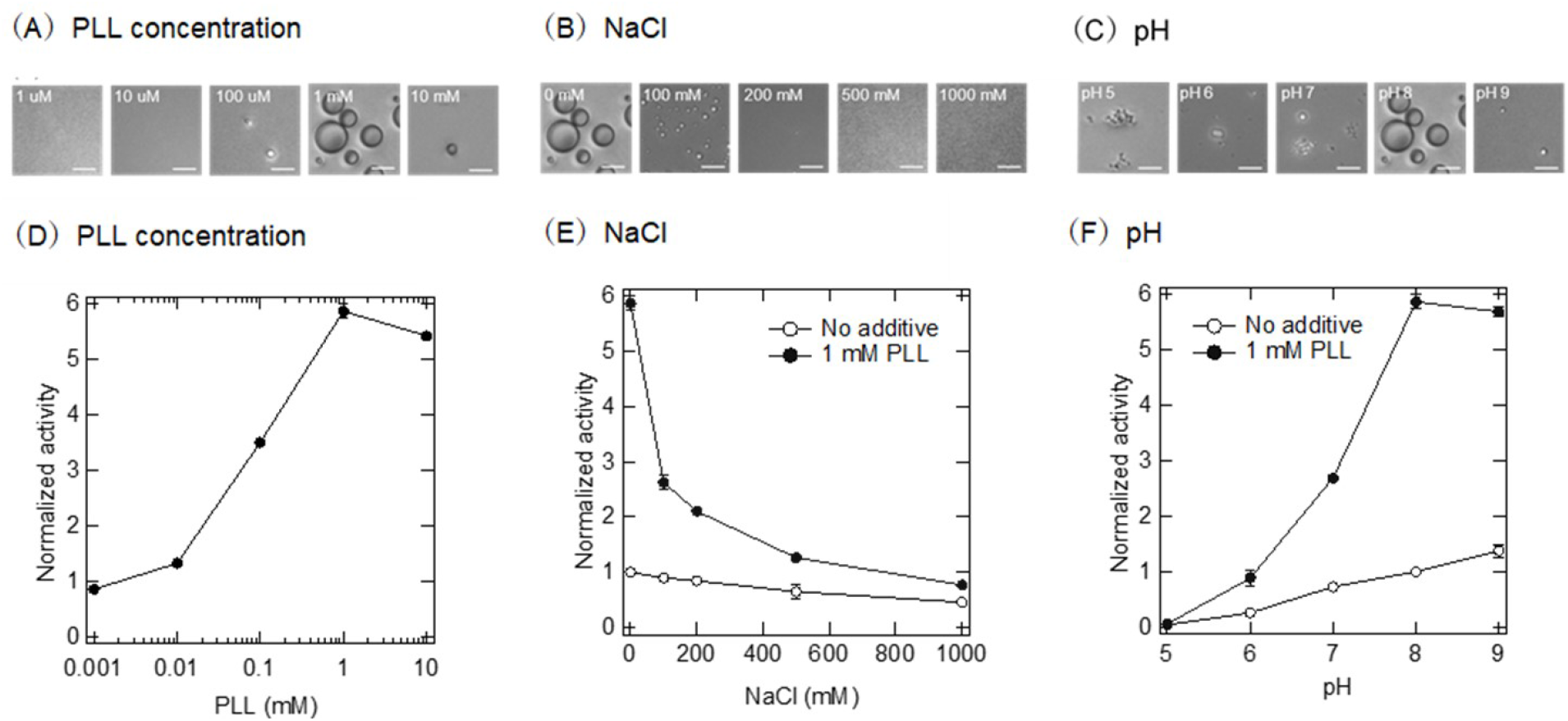
Activation of LOX in liquid droplets under various conditions. Bright-field microscopic images of LOX-PLL droplets depended on PLL concentration (A), NaCl concentration (B), and pH change (C). Scale bar, 10 μm. Normalized activities of LOX in the presence of LOX-PLL droplets depended on PLL concentration (D), NaCl concentration (E), and pH change (F). All solutions contained 5 μM LOX, 20 mM Tris HCl, and 20 mM MES. Additionally, 1 mM PLL (B, C) and pH 8 (A, B) were tested. The enzyme activity was normalized to the activity of the sample in the absence of PLL (D), in the absence of PLL and NaCl (E), or in the absence of PLL at pH 8 (F).

The enzyme activities of LOX were measured under each droplet formation condition. The normalized activity of LOX depended on PLL concentration (Fig. 3D), NaCl concentration (Fig. 3E), and pH change (Fig. 3F). The enzyme activity of LOX was activated a maximum of six times under the condition of droplet formation. A comparison of the enzyme activities in the presence or absence of PLL indicates that the droplet formation plays a crucial role in the activation of LOX.

### 3.3 LOX hyperactivation in submicrometer-scale liquid droplets

The enzyme activity increased in liquid droplets several tens of micrometers in diameter, but the size of these droplets is comparable to or larger than the size of many living cells. Therefore, we next confirmed whether smaller droplets also have an enzyme-activating effect from a biological perspective. Liquid droplets were observed above 200 nM LOX (Fig. 4A). Furthermore, the enzyme activity of LOX in the presence or absence of 1 mM PLL depended on the concentration of LOX (Fig. 4B). With increasing concentrations of LOX, the enzyme activity was increased regardless of the presence or absence of PLL, but the enzyme activities of LOX with 1 mM PLL were higher than those without PLL, indicating that the enzyme activity of LOX increased in the liquid droplets. It is noted that the activation effect was also observed for 20–100 nM LOX, at which the formation of droplets could not be confirmed by microscopy. Therefore, at 20 nM LOX, we investigated the size of the liquid droplets under various conditions by dynamic light scattering. The particle sizes of LOX-PLL droplets were dependent on PLL concentration, pH, and NaCl concentrations (Fig. 4C–E). Almost all of the LOX-PLL droplets formed average sizes from 200 nm to 1,000 nm, depending on the solution conditions. The main driving force of the formation of liquid droplets was the electrostatic interaction between LOX and PLL (Fig. 4E). Furthermore, the normalized activities of LOX depended on PLL concentration, pH, and NaCl concentrations (Fig. 4F–H). The activities of LOX shown in Figures 4F–H were similar to those of the droplet sizes shown in Figures 4C–E, indicating that the formation of submicrometer-scale liquid droplets also plays an important role in the activation of LOX.

**Figure 4.**
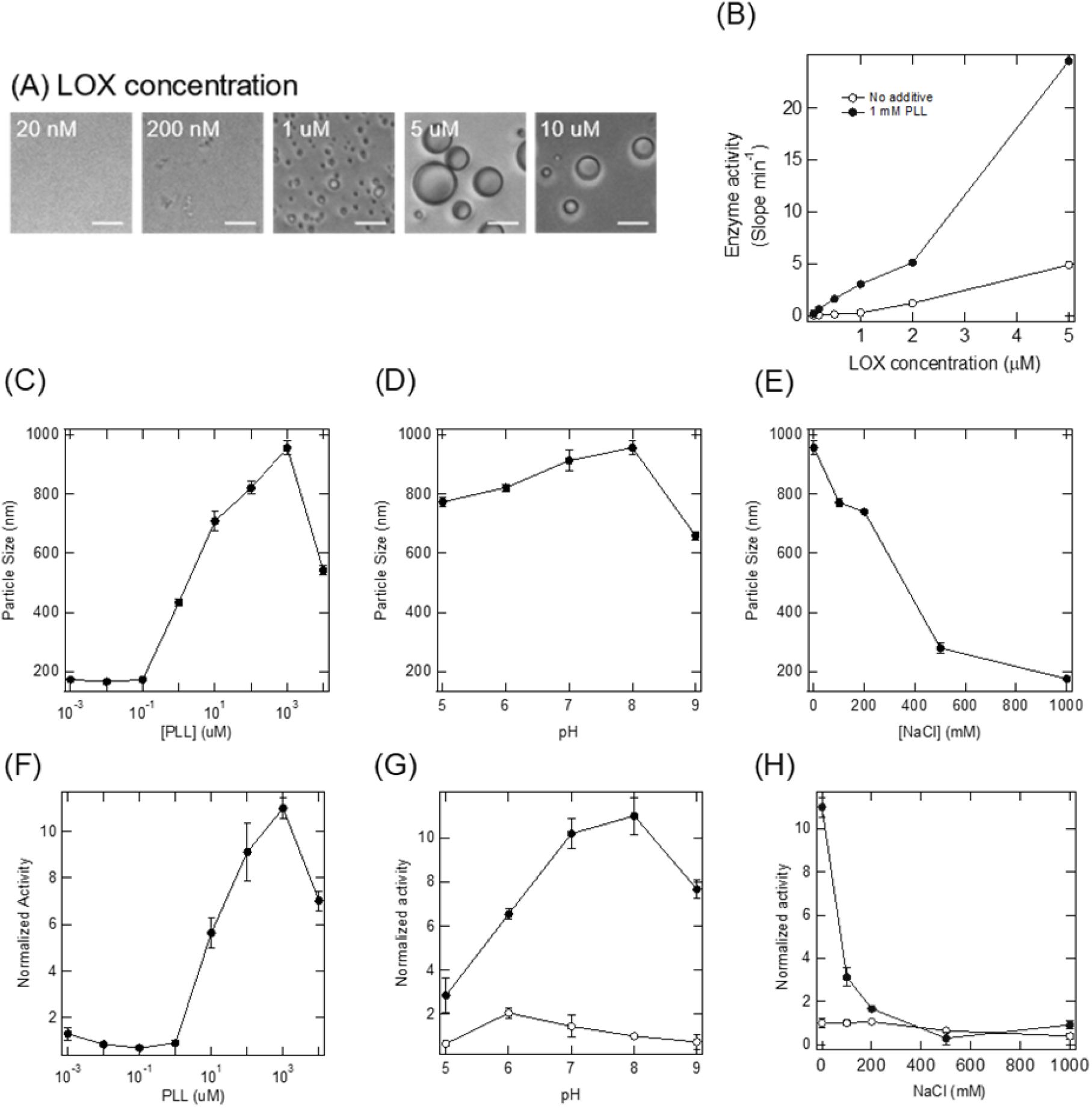
Enzyme activity of LOX in various sizes of liquid droplets (A) Microscopic images of LOX-PLL droplets depended on LOX concentration. Scale bar, 10 μm. (B) The enzyme activity of LOX depended on LOX concentration. Particle sizes of LOX-PLL droplets depended on PLL concentration (C), pH change (D), and NaCl concentration (E). The normalized activity of LOX in the presence of LOX-PLL droplets depended on PLL concentration (F), pH change (G), and NaCl concentration (H). The enzyme activity was normalized to the activity of the sample in the absence of PLL (F), in the absence of PLL at pH 8 (G), and in the absence of PLL and NaCl (H). Closed circles, activity in the presence of PLL; open circles, activity in the absence of PLL (G, H). The concentration of LOX was 20 nM (C–H).

### 3.4 Kinetic analysis of LOX in the droplets

To elucidate the detailed mechanism of LOX activation in the LOX-PLL droplets, we determined the enzyme kinetic parameters of 20 nM LOX in 20 mM Tris HCl and 20 mM MES buffer (pH 8.0) at 25°C (Fig. 5 and Table 1). The *K*_M_ of LOX with 1 mM PLL was 0.45 mM, approximately fourfold smaller than that without PLL, indicating that the liquid droplet is favorable for binding between LOX and its substrate. Additionally, the *k*_cat_ of LOX in the LOX-PLL droplet was approximately tenfold higher than that without liquid droplet, indicating that the turnover number of LOX increased in the liquid droplets. These results indicate that the synergistic effect of both substrate affinity and catalytic turnover activate LOX in the liquid droplets.

**Figure 5.**
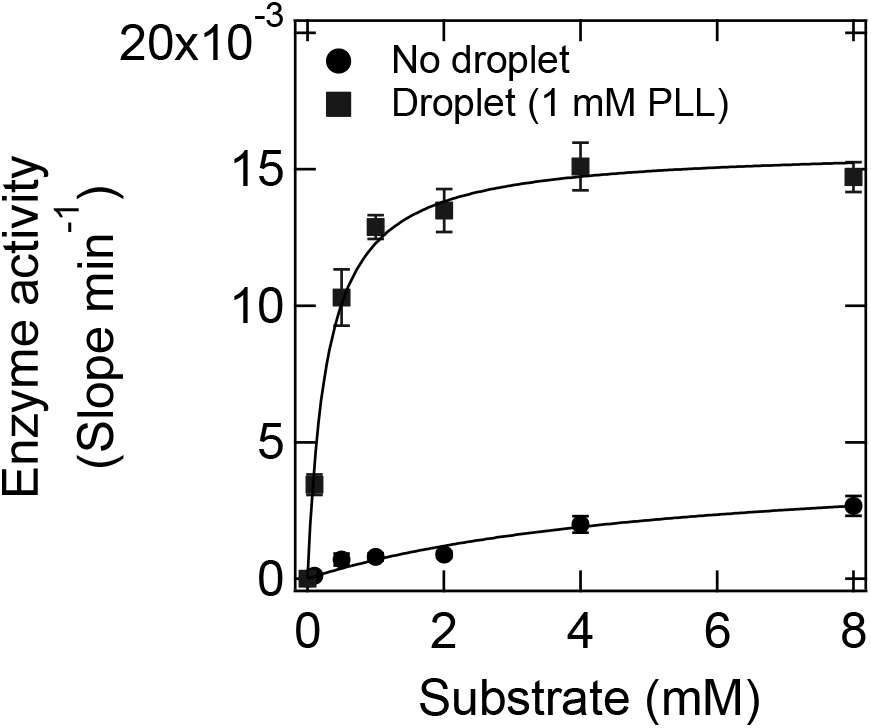
Enzyme kinetics of LOX in the droplets. The sample solution contained 20 nM LOX, 0 or 1 mM PLL, 0-8 mM L-lactic acid, 20 mM Tris-HCl, and 20 mM MES (pH 8).

**Table 1.** Michaelis-Menten parameters obtained from nonlinear fitting to the Michaelis-Menten equation

| PLL (mM) | $k_{\text{cat}}$ ($\text{s}^{-1}$) | $K_{\text{M}}$ (mM) | $k_{\text{cat}} / K_{\text{M}}$ |
| --- | --- | --- | --- |
| 0 | $71 \pm 5$ | $1.93 \pm 0.39$ | 37 |
| 1 | $638 \pm 14$ | $0.45 \pm 0.04$ | 1418 |

### 3.5 The interaction between substrate and PLL

Polymers with opposite charges to the substrate reduce *K*_M_ without the formation of liquid droplets.^19^ Because the substrate L-lactic acid is negatively charged, and PLL is positively charged, the interaction between L-lactic acid and PLL may contribute to the decrease in *K*_M_ in addition to the formation of liquid droplets. We therefore investigated the interaction of PLL and L-lactate by isothermal titration calorimetry (ITC) and NMR. The titration of L-lactate into the PLL solution showed no significant changes during the titration (Fig. 6A), indicating that L-lactate did not bind to PLL. By contrast, the titration of LOX into PLL solution showed a large positive value with a maximum enthalpy of 0.7 kcal/mol (Fig. 6B). This was due to the entropy-driven behavior observed during the formation of polymer complexes by electrostatic interactions.^20^ For confirmation of the binding between L-lactate and PLL, ^1^H-NMR spectrum of L-lactate was performed in the presence or absence of PLL (Fig. 6C). L-lactate alone was assigned by three peaks ranging 1.22–1.24, 1.34–1.38, and 1.48–1.50 ppm. The addition of PLL to the L-lactate solution did not change the intensities or the chemical shifts of these peaks. These results also indicate that L-lactate does not interact with PLL. Thus, liquid droplets may play an essential role in increasing the affinity between the substrate and enzyme, rather than between the substrate and the PLL scaffold.

**Figure 6.**
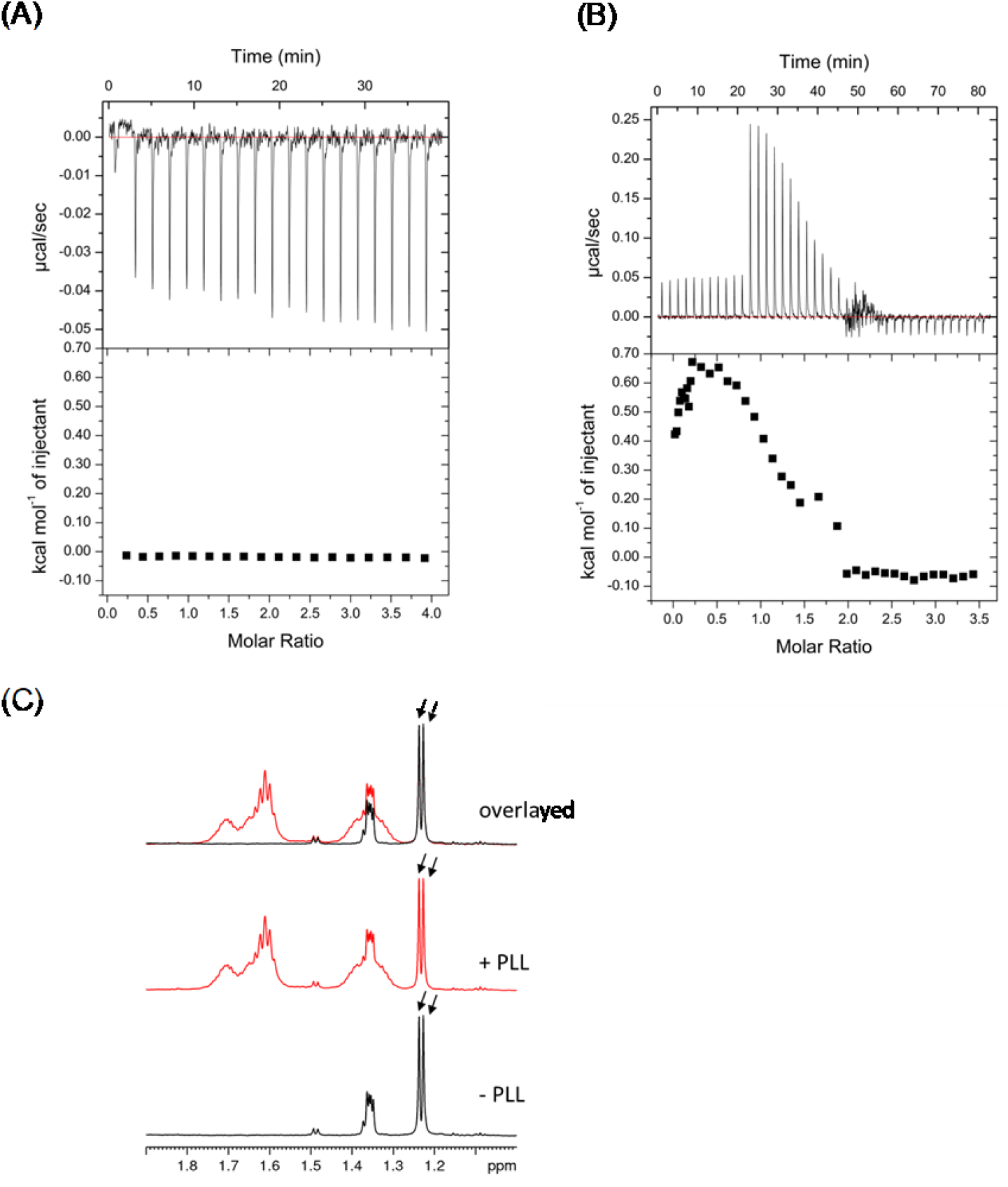
The interaction of PLL with L-lactate. ITC analysis to quantify the interaction of PLL with L-lactate (A) and LOX (B). (C) 1H-NMR spectra of PLL, L-lactate in the presence and absence of PLL, overlaid.

### 3.6 Secondary structure of LOX in the liquid droplets

Because the increase in *k*_cat_ may be due to the conformational change of LOX within the liquid droplets, we next investigated the conformational change of LOX in the liquid droplet by examining the secondary structure of LOX via the far-ultraviolet (UV) circular dichroism (CD) spectrum. The far-UV CD spectra of LOX showed negative peaks at 208 and 218 nm, and that of PLL showed positive absorption peaks at 210–230 nm (Fig. 7A). Far-UV CD spectra of the mixture of LOX and PLL did not match that calculated from the individual spectra of LOX and PLL alone (Fig. 7B). These results suggest that some of the structural changes of LOX or PLL were induced by their interaction.

**Figure 7.**
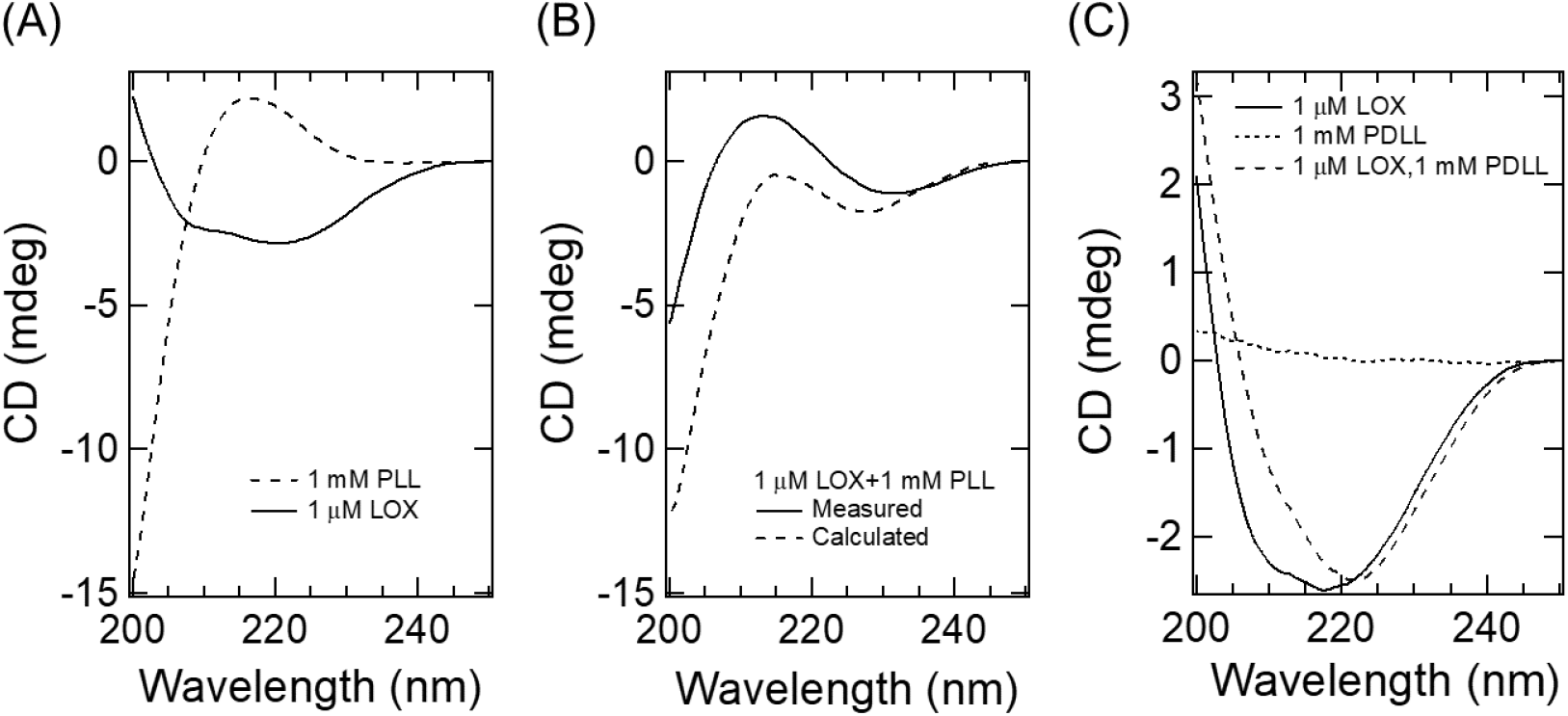
Far-UV CD spectra of LOX and PLL. (A) CD spectra of 1 μM LOX (solid line) and 1 mM PLL (broken line). (B) CD spectra of LOX and PLL mixture (solid line), and CD spectrum calculated from that of LOX and PLL in A (broken line). (C) Far-UV CD spectra of LOX and that with PDLL.

PLL or LOX alone showed large far-UV CD ellipticity (Fig. 7A). Thus, we employed poly-(D,L)-lysine (PDLL), which is achiral in CD measurements, to detect subtle changes in the secondary structure of LOX (Fig. 7C). As expected, PDLL showed almost no far-UV CD spectrum due to its lack of optical activity (Fig. 7C). In the presence of 1 mM PDLL, at which LOX and PDLL formed a liquid droplet and showed hyperactivation (Supplementary Fig. S1), the spectrum of LOX drastically changed. Notably, the intensity of LOX at 210 nm was higher than that in the absence of PDLL. These results indicate that a conformational change of LOX was induced in the liquid droplet, which may cause its increased *k*_cat_.

## 4. Discussion and conclusion

Here, we reported that the enzyme activity of LOX in liquid droplets was extremely enhanced compared with that in the dispersion state. The LOX-PLL droplets were mainly stabilized by electrostatic interactions between anionic LOX and cationic PLL (Figs. 3 and 4). The mechanism of LOX enzyme activation might be derived from the kinetic parameters (Fig. 5 and Table 1) and conformational properties (Fig. 7). Briefly, the formation of liquid droplets decreased *K*_M_ due to the compartmentalization of substrates in the liquid droplet, not due to the interaction between substrate and PLL (Fig. 6). Interestingly, the liquid droplets increased *k*_cat_ about tenfold higher than the buffered solution. To our knowledge, this is the first report of increased *k*_cat_ in liquid droplets. The activation of LOX in the liquid droplets resulted from its conformational change, as suggested by the CD results (Fig. 7).

Enzyme activation by the formation of liquid droplets may result from one or more of the following possibilities. First, the interaction of the enzyme with the polymer in the liquid droplets may increase the structural stability of the native enzyme, leading to an increased *k*_cat_.^21–23^ Second, liquid droplets are highly crowded with macromolecules, resulting in the exclusion of water molecules. This crowding stabilizes nonnative structures that differ from those in dilute conditions.^24^ Therefore, the crowding effect may promote the transition state of the enzyme, leading to the enhancement of enzyme activity. Finally, liquid droplets represent a nonpolar environment compared with a buffered solution.^25^ A nonpolar solution increases the stability of hydrogen bonds and electrostatic interactions, which influence enzyme activity. Furthermore, LOX has a disordered loop and a short helix that control enzyme activity.^26,27^ These mobile regions cover the active center, and hence the flexibility of the regions contribute to the activation enzyme, such as substrate uptake, product release, and the formation of an active site microenvironment. Specifically, the hydrogen bond network in the active site and product pyruvate play important roles in the activity of LOX. The CD results show that the percentage of LOX α-helix within the droplet was reduced, indicating changes in the structure and dynamics of this region within the droplet. Further structural analysis of this region within droplets will be an interesting subject for future research.

The activation of LOX within liquid droplets is valuable information for industrial applications. Currently, the primary method of enzyme activation is protein engineering^28^ and directed evolution.^29^ These methods create favorable mutants with high activity by repeated mutation introduction and screening. However, they are time-consuming and costly to produce desirable mutants. By contrast, the formation of liquid droplets is a simple method for improving enzyme activity because only the addition of a poly-electrolyte into the enzyme solution is needed. LOX is an oxidoreductase applied to biofuel cells^18^ and biosensors.^30^ Thus, the formation of liquid droplets represents a versatile approach for improving enzyme activity for practical applications using LOX or other enzymes.

## ACKNOWLEDGMENT

This work was partially supported by JST-ASTEP (Grant No. AS272S004a) to T.M.. This work was partly supported by JSPS KAKENHI (Grant No. 18H02383 and 18H01719) to K.S..

