## Supplemental Figure 1 for "Hyperactivation of L-lactate oxidase by liquid-liquid phase separation"

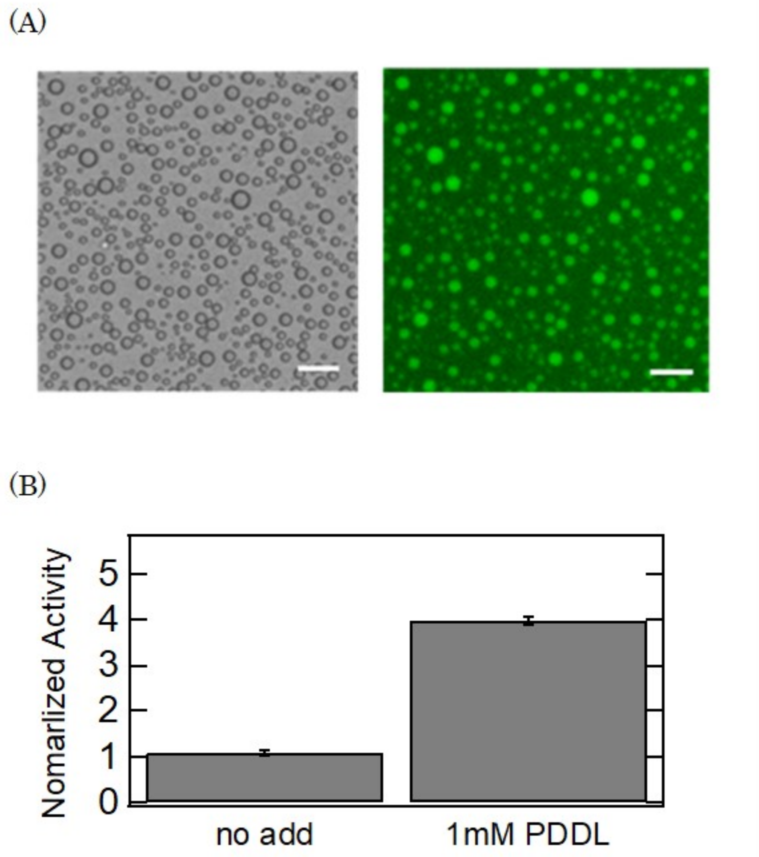


Figure S1 Formation of LOX -PDDL droplets and activation.
(A) Blight field microscopic images of droplet (Left) and fluorescent microscopic images of LOX (Right) The solution contained 5 μM LOX, 1 mM PDLL, 20 mM Tris HCl, and 20 mM MES (pH 8). Scale bar, 20 μm.
(B) Normalized activity of LOX in the presence of 1 mM PDLL.
